# Nuclear actin interactome analysis links actin to KAT14 histone acetyl transferase and mRNA splicing

**DOI:** 10.1101/445973

**Authors:** Tiina Viita, Salla Kyheröinen, Bina Prajapati, Jori Virtanen, Markku Varjosalo, Maria K. Vartiainen

## Abstract

In addition to its essential functions within the cytoskeleton, actin also localizes to the cell nucleus, where it is linked to many important nuclear processes from gene expression to maintenance of genomic integrity. However, the molecular mechanisms by which actin operates in the nucleus remain poorly understood. Here we have used two complementary mass spectrometry (MS) techniques, AP-MS and BioID-MS, to identify binding partners for nuclear actin. Common high-confidence interactions highlight the role of actin in chromatin remodeling complexes and identify the hATAC histone modifier as a novel actin-containing nuclear complex. Further analysis demonstrates that actin binds directly to the hATAC subunit KAT14, and modulates its histone acetyl transferase activity *in vitro* and in cells. BioID-MS, which can detect also transient interactions, links actin to several steps of transcription as well as to RNA processing. Alterations in nuclear actin levels disturb alternative exon skipping of the SMN2 minigene, suggesting also a functional role for actin in RNA splicing. This interactome analysis thus identifies both novel direct binding partners and functional roles for nuclear actin, as well as forms a platform for further mechanistic studies on how actin operates during essential nuclear processes.

## Introduction

Actin and its importance in the cytoplasm is well known. Actin filaments and their coordinated assembly, regulated by numerous actin-binding proteins (ABPs), are required for cell migration, cytokinesis and membrane dynamics (Le Clainche and Carlier, 2008; Pollard and Cooper, 2009). Although actin is more recognized from its actions in the cytoplasm, it was found already decades ago inside the nucleus and suggested to be involved in transcription (Egly et al., 1984). Today, actin is linked to numerous nuclear functions from gene expression to maintenance of genomic integrity (Viita and Vartiainen, 2017).

Development of probes, which recognize different forms of actin, as well as improved live imaging techniques, have proven the significance of actin dynamics in the nucleus (Grosse and Vartiainen, 2013). Nowadays it is firmly established that nuclear actin can polymerize, and even form phalloidin-stainable filaments in certain situations, such as upon serum stimulation (Baarlink et al., 2013), cell spreading (Plessner et al., 2015), DNA damage responses (Belin et al., 2015; Schrank et al., 2018; Wang et al., 2017) and during certain cell-cycle phases (Baarlink et al., 2017). In the cytoplasm, a large number of actin-binding proteins regulate actin dynamics. However, despite the fact that many actin-binding proteins localize to the nucleus (Kumeta et al., 2012), only very few of them have been shown to regulate nuclear actin dynamics. These include cofilin (Baarlink et al., 2017), diaphanous-related formins (mDia1/2) (Baarlink et al., 2013), formin-2 (FMN2), together with Spire-1/Spire-2 (Belin et al., 2015) and Arp2/3 complex (Caridi et al., 2018; Schrank et al., 2018). Collectively these studies have demonstrated that like cytoplasmic actin, nuclear actin dynamics are very tightly regulated.

Actin is linked to many processes that regulate gene expression. Actin, often together with the actin-related proteins (Arps), is a component of many chromatin remodeling complexes such as SWI/SNF (Zhao et al., 1998), SWR1 (Mizuguchi et al., 2004), Nu4A (Galarneau et al., 2000), Ino80 (Ayala et al., 2018; Eustermann et al., 2018), Tip60 (Ikura et al., 2000) and SRCAP (Cai et al., 2005). It has been shown that different ATPases of the chromatin remodeling complexes can bind actin with their helicase-SAINT associated (HSA) domain (Szerlong et al., 2008). Recent structural work have begun the shed light on the functional relevance of actin in these complexes. For example, in the Ino80 complex actin, Arp4 and Arp8 form a module that is involved in recognizing the extranucleosomal linker DNA (Knoll et al., 2018). Actin can also regulate gene expression by controlling the activity of specific transcription factors. Perhaps the best-characterized example is serum response factor (SRF), which controls the expression of many cytoskeletal genes in response to changes in actin dynamics. The signal from the actin cytoskeleton to SRF is mediated by the transcription coactivator Mrtf-A (also known as MAL or MKL1) (Miralles et al., 2003). Mrtf-A binds actin monomers through its RPEL domain, which regulates the nuclear localization and activity of Mrtf-A in response to actin dynamics (Vartiainen et al., 2007).

Actin has also been linked directly to the transcription process, because it can be co-purified with all three RNA polymerases (Pol): Pol I (Fomproix and Percipalle, 2004), Pol II (Egly et al., 1984; Smith et al., 1979) and Pol III (Hu et al., 2004). The precise molecular mechanisms by which actin participates in transcription are still unclear, but balanced nucleo-cytoplasmic shuttling of actin is necessary for transcription (Dopie et al., 2012; Sokolova et al., 2018). Availability of nuclear actin monomers seems to be critical here, since polymerization of nuclear actin to stable filaments (Serebryannyy et al., 2016b) or activation of a mechanosensory complex consisting of emerin, non muscle myosin II and actin (Le, et al 2016) repress transcription. Our recent genome-wide analysis revealed that actin interacts with essentially all transcribed genes. Actin is found, together with Pol II, near transcription start sites of most genes, as well as on the gene bodies of highly expressed genes (Sokolova et al., 2018). Actin may thus have several functions during the transcription process, and has, in fact, been implicated in pre-initiation complex (PIC) assembly (Hofmann et al., 2004) as well as transcription elongation via pTEF-β complex (Qi et al., 2011). In addition, actin has also been shown to bind heterogeneous nuclear ribonucleoproteins (hnRNPs) (Obrdlik et al., 2008; Percipalle et al., 2003; Percipalle et al., 2002), which could indicate a function for actin in mRNA processing.

Beyond gene expression, nuclear actin has recently been linked to also DNA damage responses and DNA replication. Actin dynamics and formin activity are required for initiation of DNA replication by influencing nuclear transport and loading of replication proteins onto chromatin (Parisis et al., 2017). Nuclear actin dynamics are also important in DNA damage responses as reduced nuclear actin increases the number of double stranded DNA breaks (DSB) in cells (Belin et al., 2015). Moreover, it seems that DNA damage promotes nuclear actin filament formation (Belin et al., 2015; Wang et al., 2017). These filaments are needed for the DNA damage repair, as their loss leads to reduced efficiency of DSB clearance. Two recent papers show that the nuclear actin polymerization, mediated by the Arp2/3 complex, is needed for DBS movement in the nucleus (Caridi et al., 2018; Schrank et al., 2018).

It is evident that actin has many important functions within the nucleus. However, the molecular mechanisms by which actin operates during these essential events have remained largely unclear, with one critical aspect being the relatively limited knowledge that we have on the binding partners for nuclear actin. For this reason, we decided to resolve the nuclear actin interactome by using two complementary mass spectrometry (MS)-based techniques, Affinity Purification (AP)-MS and Proximity-dependent biotin identification (BioID)-MS, allowing us to probe both stable and dynamic interactions of nuclear actin. As expected, the stable interactions include several components of different chromatin remodeling complexes known to contain actin. The BioID data, on the other hand, links actin to pre-mRNA processing and transcription. Among the hits, we identify a novel direct binding partner for actin in the nucleus, KAT14, which is part of the histone modifying complex Ada-Two-A-Containing complex (hATAC). Further functional analysis reveals that actin inhibits the KAT14 histone transferase activity both *in vitro* and in cells. Collectively, our nuclear actin interactome analysis links actin to novel functions within the nucleus, and provides a platform for further mechanistic studies.

## Results

### Two complementary approaches for obtaining the nuclear actin interactome

Actin has been linked to many important nuclear processes from gene expression to maintenance of genomic integrity, but the molecular mechanisms have remained largely unclear. The first step towards elucidating these mechanisms is to identify the binding partners for actin in the nuclear compartment. Here we have used two complementary Mass Spectrometry (MS) based approaches, Affinity Purification (AP)-MS (Varjosalo et al., 2013) and Proximity-dependent biotin identification (BioID)-MS (Roux et al., 2012) (Figure 1A) to identify binding partners for nuclear actin. These two methods complement each other, because AP-MS aims to catch stable protein complexes in mild lysis conditions, while biotinylation of near neighbors of the bait protein in BioID allows the detection of more transient interactions, and the usage of harsher lysis conditions. The low amount of actin in the nucleus compared to cytoplasm, and the ability to distinguish the actual nuclear interactions from cytoplasmic ones [reviewed in (Viita and Vartiainen, 2017)], make it challenging to study nuclear actin-binding partners. To overcome these obstacles in our study, we have added nuclear localization signal (NLS) to actin to enrich actin amounts in the nucleus. As a control, besides of the diffusively localizing GFP (Supplementary Figure 1C), we used actin without the NLS, which allowed us to compare cytoplasmic and nuclear pools of actin with our two MS techniques (Supplementary Figure 1A,B; see Materials and methods for MS data analysis). Primary data from the MS can be found in Supplementary table 1 and analyzed data with high-confidence interactions in Supplementary table 2. Database for Annotation, Visualization and Integrated Discovery (DAVID) (Huang et al., 2009) annotation tool revealed that NLS-signal increases the amount of proteins with GO term nucleus (GO: 0005634) (58 % in AP-MS-NLS-actin and 65 % in BioID-NLS-actin) compared to the interactomes without NLS-signal (38 % in AP-MS-actin and 0 % in BioID-actin) (Figure 1B). This shows that we have been able to enrich nuclear proteins with our NLS-actin constructs. We have also used non-polymerizable R62D-actin mutant (Supplementary Figure 1 A,B) (Posern et al., 2002) to study if polymerization status of actin would affect its nuclear interactions. However, most NLS-Actin and NLS-R62D-Actin hits were overlapping (Figure 1C), which indicates that actin does not need the capacity to polymerize to interact with the proteins detected in our experimental setup. For this reason, we combined the actin and actin-R62D datasets and thus overall obtained 4 different interactomes: NLS-actin AP-MS, NLS-actin BioID, actin AP-MS and actin BioID (Supplementary table 2). The shared hits from the actin AP-MS and BioID were, as expected, known regulators of the cytoskeleton and of actin filament assembly (Supplementary Figure 2 A,B).

**Figure 1.**
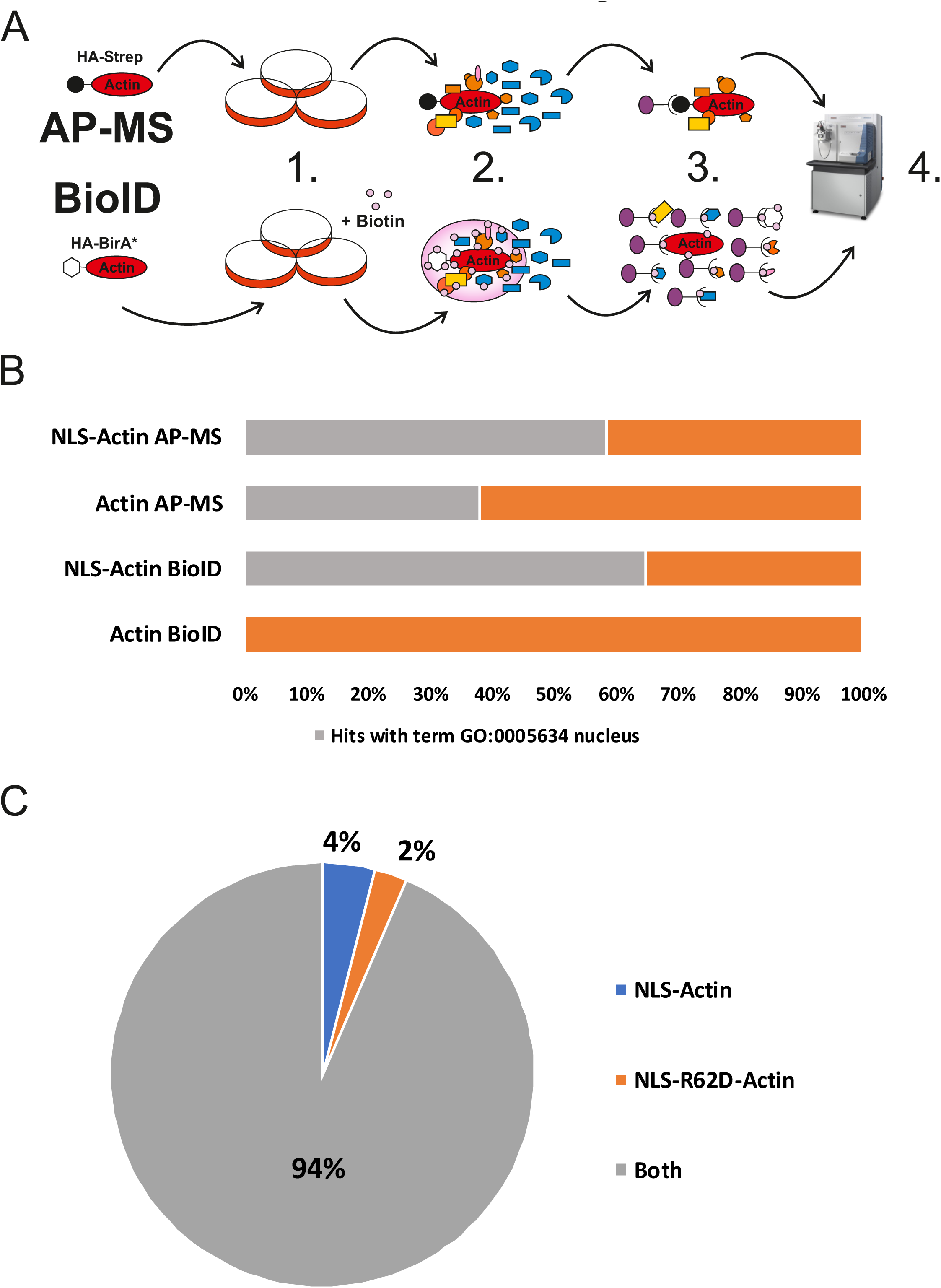
Nuclear actin interactome resolved by using two different MS-approaches and differently localizing actin constructs. A. Schematic view of two different MS-approaches, Affinity Purification MS (AP-MS) and Proximity-dependent biotin identification (BioID). 1. Stable, inducible HEK Flp-In cell lines expressing different actin constructs tagged with HA-Strep (hemagglutinin Strep-tag) or HA-BirA* (mutated minimal biotin ligase) 2. After induction of actin construct expression, protein complexes are formed with tagged proteins. In BioID, addition of biotin allows HA-BirA* to biotinylate near neighbors of the tagged protein. 3. Single step purification with StrepTactin to purify formed protein complexes. Milder lysis conditions used in AP-MS to obtain full, intact protein complexes. Purification of biotinylated proteins in BioID-MS enables usage of harsher lysis conditions. 4. Protein digestion to peptides and LC/MS analysis. B. Fraction of high-confidence interactions with the Cellular Component (CC) GO term nucleus (GO:0005634) for the different actin constructs. C. Pie chart showing the fraction of hits that are unique or shared for NLS-actin and NLS-R62D-actin from both AP-MS and BioID.

### Stable interactions of actin in chromatin remodeling and modifying complexes

AP-MS resulted in fewer high-confidence interactions than BioID (Figure 2A), likely because BioID can also detect more transient interactions. Similar differences between these two approaches have been observed previously for various proteins (Lambert et al., 2015; Liu et al., 2018). Most of the shared hits from the AP-MS and BioID (Figure 2B) include already established interactions of actin in chromatin remodeling complexes, including SWI/SNF (Zhao et al., 1998) (ARP4, BRG1 and BAF170) and SRCAP/TIP60 (Cai et al., 2005; Ikura et al., 2000) (ARP4, EP400, YEATS4, DMAP1). Three shared hits, KHSRP (Russo et al., 2011), hnRNPF (Wang and Cambi, 2009) and SSB (Liang et al., 2013), have been linked to alternative splicing. In addition, the shared hits contained several subunits (KAT14, ZZZ3, MBIP, YEATS2) of the human Ada-Two-A-containing (hATAC) complex (Guelman et al., 2009; Wang et al., 2008). To our knowledge, hATAC has not been linked to actin before, although the complex interacts with PCAF, which in turn binds actin (Obrdlik et al., 2008). hATAC complex is a histone acetyl transferase complex, which can acetylate H3 and H4 in mammals (Guelman et al., 2009) and in fruit fly (Guelman et al., 2006; Suganuma et al., 2008).

**Figure 2.**
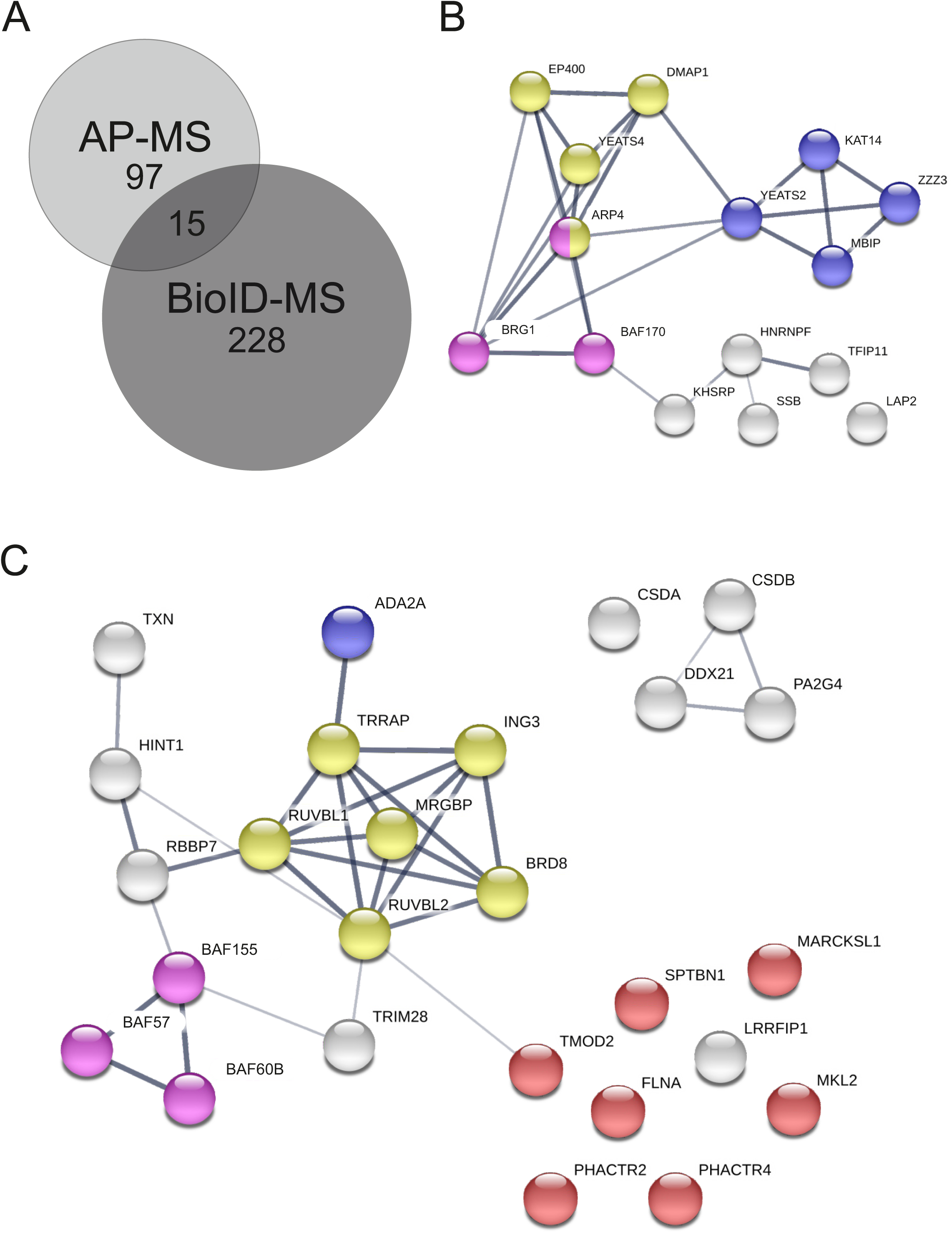
Shared hits from AP-MS and BioID link actin mainly to chromatin remodeling and modifying complexes. A. Number of unique and shared high confidence interactions from AP-MS and BioID for nuclear actin (combined NLS-actin and NLS-actin-R62D). B. Shared hits from nuclear actin AP-MS and BioID shown as a STRING map. SWI/SNF complex (GO:0016514) highlighted in pink, ATAC complex (GO:0005671) in blue, and Nu4A histone acetyltransferase complex (GO:0035267) (Nu4A is the core complex of SRCAP/Tip60) in yellow. C. High-confidence interactions for nuclear actin obtained only with the AP-MS technique with Swiss-Prot keywords actin-binding, chromatin remodeling and transcription (26 out of total 96 hits) shown as a STRING map. Different complexes are highlighted as in B, with actin-binding proteins in red.

The unique hits from the AP-MS contained further subunits from the SWI/SNF (BAF155, BAF57 and BAF60b) (Euskirchen et al., 2012), SRCAP/TIP60 (RUVBL1/2, TRRAP, BRD8, MRGBP and ING3) (Doyon et al., 2004) and hATAC (ADA2A) (Guelman et al., 2009; Wang et al., 2008) complexes (Figure 2C), which shows that we were able to pull down intact complexes with this technique. In addition, these hits also contained several proteins, which are known to interact with actin, including PHACTR2, PHACTR4 (Allen et al., 2004), FLNA (Shao et al., 2016), TMOD2 (Fowler and Dominguez, 2017), MARCKSL1 (El Amri et al., 2018), SPTBN1 (Machnicka et al., 2014) and MRTF-B (Miralles et al., 2003) (Figure 2B). Many of these proteins, or their homologs, have also previously been localized to the nucleus (Deng et al., 2012; Machnicka et al., 2014; Miralles et al., 2003; Rohrbach et al., 2015; Wiezlak et al., 2012). For instance, MRTF-B is a transcription coactivator of SRF, and presumably regulated by nuclear actin similarly to MRTF-A (Vartiainen et al., 2007). FLNA, an actin-crosslinking protein, interacts with MRTF-A to regulate its activity (Kircher et al., 2015) and has also been linked to DNA repair (Yue et al., 2009), whereas LRRFIP2 is an transcription coactivator (Liu et al., 2005) that interacts with Flightless-I, an actin-binding protein of the gelsolin family (Fong and de Couet, 1999). Finally, over 30% of the unique hits from the AP-MS were linked to translation (Supplementary table 2). This may reflect the fact that actin has been linked to both Pol I-mediated transcription and assembly of ribonucleoprotein particles (Percipalle, 2009).

### BioID links actin to transcription and mRNA processing

The high-confidence interactions from BioID link actin to many different functions in the nucleus such as chromatin remodeling, transcription, DNA replication and mRNA processing (Figure 3A). Most notably, more than 30 % of the unique hits from BioID, as well as few unique hits from the AP-MS (DDX21, CSDB), were associated with mRNA splicing or processing (Figures 2B and 3A). The hits included small nuclear ribonucleoproteins (snRNPs) from essentially all of the major subunits of the spliceosome and proteins regulating different spliceosome compositions, particularly those of complex B or B^act^ (Figure 3B) (Wahl and Luhrmann, 2015). These findings imply that actin could be directly involved in pre-mRNA splicing/processing. Alternatively, the co-transcriptional nature of splicing (Herzel et al., 2017), and the involvement of actin in transcription (Fomproix and Percipalle, 2004; Hofmann et al., 2004; Kukalev et al., 2005; Obrdlik et al., 2008; Percipalle et al., 2003; Philimonenko et al., 2004; Sokolova et al., 2018), may result in the detection of these RNA splicing/processing related proteins here.

**Figure 3.**
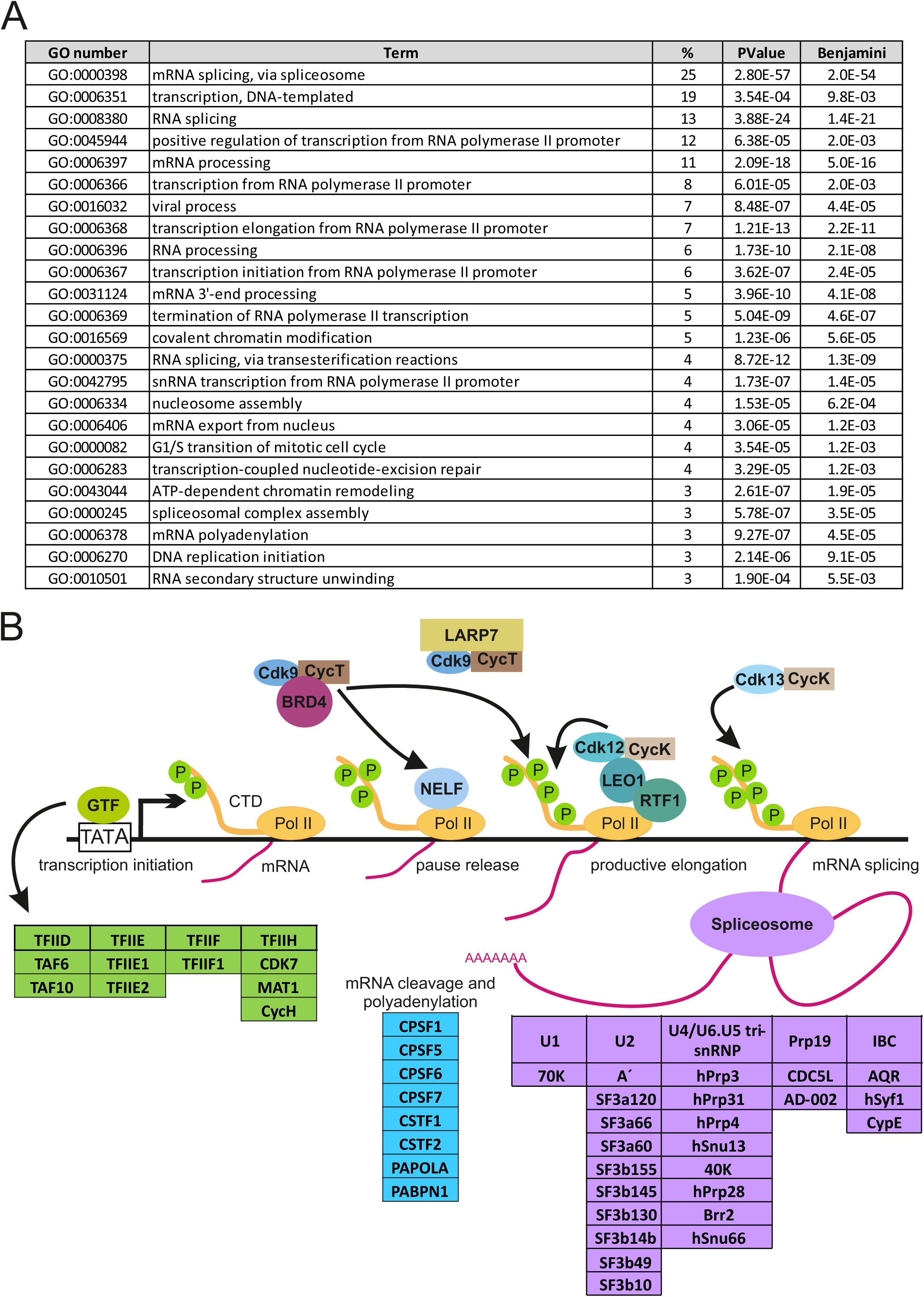
BioID links actin to mRNA processing and transcription. A. DAVID functional annotation chart (Cut off Benjamini 10^−3^ and >2 % of all hits) with GO Direct Biological Pathways from unique BioID hits. B. Schematic picture of high-confidence interactions from BioID for nuclear actin that are involved in different stages of transcription (AJ et al., 2016; Gruber et al., 2014; Wahl and Luhrmann, 2015; Van Oss et al., 2017). Compositions of general transcription factors (GTFs) were obtained from KEGG pathway Ko03022 (Kanehisa and Goto, 2000). Note that RNA polymerase II (Pol II) is included for visualization purposes, although it was not a hit in the MS screens for nuclear actin. Phosphatase group is indicated with green circle with P and arrows indicate regulatory functions.

Indeed, the unique hits from BioID link actin to different stages of transcription from the assembly of the PIC to transcription elongation by Pol II (Figure 3B). The hits included for example subunits of the TFIID, TFIIE, TFIIF and TIIH general transcription factors, supporting the previous data that link actin to PIC formation (Hofmann et al., 2004). In terms of transcription elongation, the pTEF-β complex subunits, CDK9 and Cyclin T, which have also previously been linked to monomeric actin (Qi et al., 2011), were hits in our BioID screen (Figure 3B, Supplementary table 2). Interestingly, also pTEF-β regulatory proteins LARP7 (BioID), BRD4 (BioID) and DDX21 (AP-MS) (AJ et al., 2016) were identified as putative interactors for nuclear actin, indicating that actin could regulate the activity of pTEF-β by modulating its association or release from these factors. Curiously, the BioID hits included also CDK12, CDK13 and Cyclin K, which have all been implicated in transcription elongation (Greifenberg et al., 2016). Further experiments are needed to elucidate, if actin generally regulates CDK-mediated Pol II CTD phosphorylation like it has been postulated for the actin-CDK9 interaction (Qi et al., 2011). Of note, none of the RNA polymerase subunits were identified as high-confidence interactions in our assays (Supplementary table 2). In agreement with earlier studies (Obrdlik et al., 2008; Percipalle et al., 2002; Percipalle et al., 2001), we detected interactions between actin and several hnRNP proteins and hnRNP-associated proteins, including hnRNPF (shared hit), hnRNPM (AP-MS), hnRNPP2 (BioID), and PSF (BioID). Recent studies have linked actin to DNA replication (Parisis et al., 2017) and our BioID screen suggested putative interactions between actin and several replication-linked proteins, including ORC2, ORC5, ORCA, POLA1, POLA2, PRIM1 and PRIM2 (Supplementary table 2) (Li and Stillman, 2012; Wu et al., 2014).

To further validate the shared hits from the AP-MS and BioID interactomes, we used light microscopy based Bimolecular Fluorescence Complementation (BiFC) technique (Cabantous et al., 2005; Kerppola, 2008; Zhou et al., 2011), which allowed us to confirm the possible interactions in intact cells (Figure 4A). For this purpose, we decided to generate stable, inducible cell line expressing HA-GFP1-10-actin and attach smaller, epitope-like tag Flag-GFP11 to our MS hits. To test our method, we used Flag-GFP11-actin construct together with our inducible HAGFP1-10-actin cell line and saw reformed GFP (BiFC signal) in transfected cells with tetracycline induction, but not in non induced cells (Figure 4B). Of the eight (BRG1, DMAP1, KAT14, MBIP, hnRNPF, SSB, TFIP11 and LAP2) tested proteins, representing different complexes and biological functions of the putative interactors, we were able to verify six by BiFC (Figure 4C). Only MBIP and LAP2 did not produce detectable GFP fluorescence, despite the fact that both were efficiently expressed in the cells (Figure 4C). This could indicate that these proteins are either false positives from the MS screens, that the interaction is not direct or that the orientation of the interaction does not allow the reformation of the GFP molecule.

**Figure 4.**
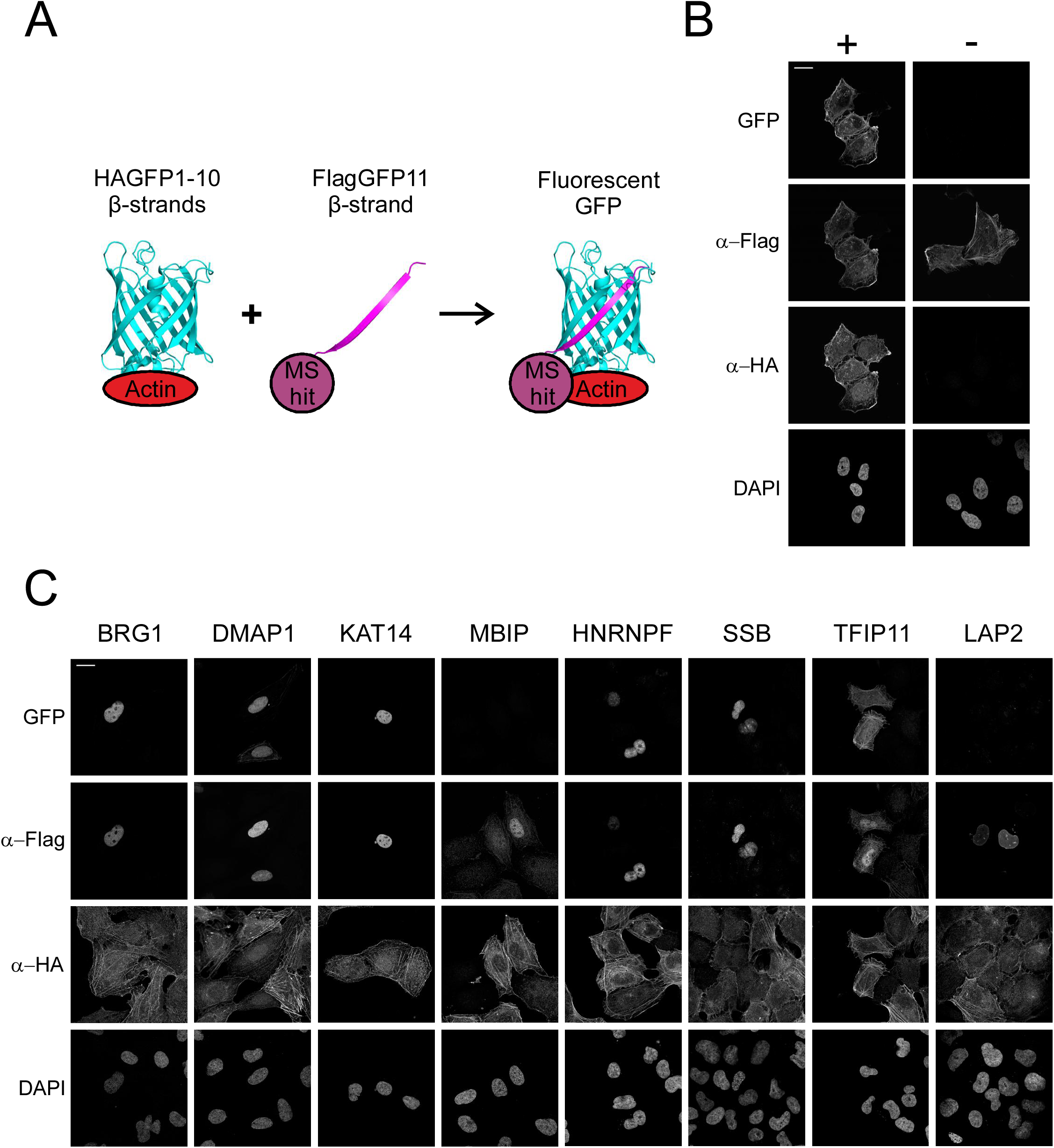
Bimolecular Fluorescence Complementation (BiFC) technique validates interactions between shared hits from the AP-MS and BioID interactomes. A. Schematic of BiFC technique, HA-GFP1-10 attached to bait protein (actin) and Flag-GFP11 attached to prey protein (MS hit). Fluorescent GFP is reformed if bait and prey interact with each other. Schematic adapted from previous publications (Cabantous et al., 2005; Zhou et al., 2011) and drawn with PyMOL and GFP reference was taken from PDB ID 5B61 (Choi et al., 2017). B. Confocal microscopy images of stable, inducible U2OS cell line expressing HAGFP1-10-actin induced (+) and non-induced (−) with tetracycline, transiently transfected with FlagGFP11-actin and stained with DAPI as well as HA- and Flag-antibodies to visualize the expression of the BiFC constructs. GFP channel shows the BiFC signal signifying the interaction between the bait and prey protein. Scale bar, 20 μm. C. BiFC microscopy assay with shared hits from the AP-MS and BioID screens. Confocal microscopy images of tetracycline induced HAGFP1-10-actin U2OS cells transiently transfected with the indicated FlagGFP11 constructs visualized as in B. Scale bar, 20 μm.

Taken together, by utilizing two complementary MS techniques we have been able to obtain a global view of the nuclear actin interactome. The identified interactions include already established binding partners for nuclear actin, as well as novel interactions that link actin to previously described functions such as chromatin remodeling, DNA replication and transcription. These novel interactions can now be used as the basis for unravelling the molecular mechanisms by which actin operates during these essential nuclear events. In this manuscript, we will further focus on the possible novel functions of actin in the context of the hATAC complex and pre-mRNA splicing.

### Actin binds directly to KAT14 histone acetyl transferase (HAT) and modulates its HAT activity

As mentioned above, hATAC is a histone acetyl transferase (HAT) complex, which consists of multiple subunits (Guelman et al., 2009; Nagy et al., 2010; Wang et al., 2008). Actin usually interacts with chromatin remodelers with nuclear Arps [reviewed in (Kapoor and Shen, 2014)], and quite unexpectedly hATAC complex does not contain nuclear Arps. This intrigued us to further investigate the relationship between actin and hATAC. ATAC complex can also interact with PCAF, which is a HAT earlier associated to actin and hnRNPU (Obrdlik et al., 2008). Interestingly, we did not obtain PCAF in our nuclear actin MS analysis (Supplementary table 2), which might hint that there is another binding partner for actin in the ATAC complex. Here we turned our attention to KAT14, since it produced very strong signal in the BiFC assay.

We first verified the interaction between KAT14 and actin by co-immunoprecipitation (Co-IP) with overexpressed proteins utilizing both proteins as baits (Figure 5A). Additionally, Co-IP with overexpressed KAT14 was able to pull down endogenous actin (Figure 5A). As our interactome data suggested that actin does not require its capacity to polymerize to associate with hATAC (Supplementary table 1), we decided to confirm this with Co-IP. We did Co-IPs with different actin mutants: non-polymerizable actin mutant (R62D-actin) and a mutant, which favors polymerization (V159N-actin) (Posern et al., 2002). In addition, we also tested an actin mutant, which has mutations in the hydrophobic cleft (G168D, Y169D-actin) (Cao et al., 2016). Quantified results from the western blots suggest that R62D-actin binds KAT14 more efficiently than wild type actin, and that the mutation in the hydrophobic cleft diminishes the interaction almost completely (Figure 5B). There was a small decrease in the signal with the polymerization favoring V159N-mutant compared to wild type actin (Figure 5C), which strengthens the hypothesis that KAT14 would prefer binding to monomeric actin. We also mapped the binding site for actin in KAT14 with N- and C-terminal truncations of KAT14 (KAT14-1-546, and KAT14-547-782) (Figure 5C). Co-IP experiments showed actin-binding to the KAT14-547-782 construct, but not to KAT14-1-546, demonstrating that the actin-binding site in KAT14 is in the C-terminus.

**Figure 5.**
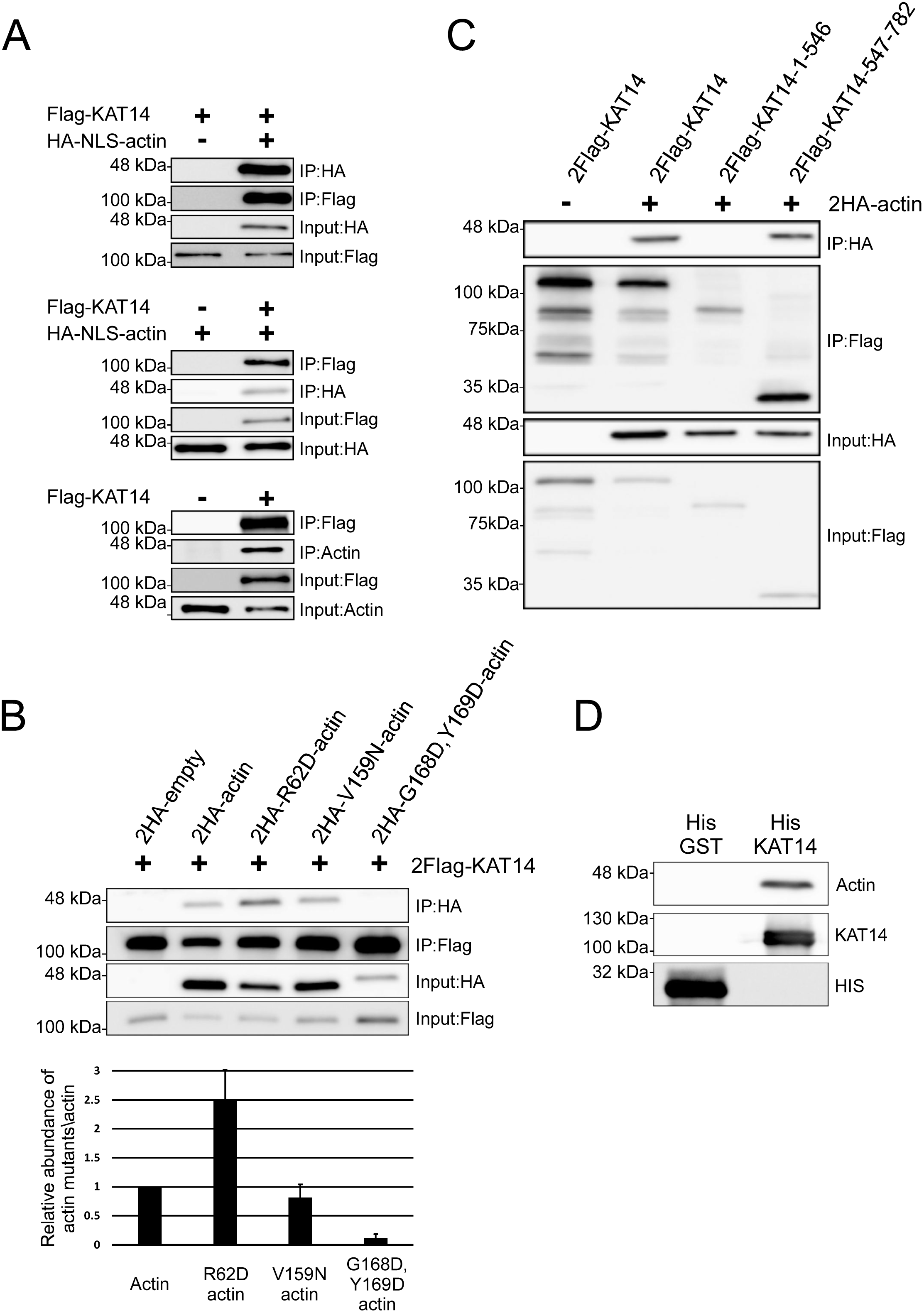
hATAC subunit KAT14 directly binds actin and favors binding to actin monomers via its C-terminus. A. Western blots of co-immunoprecipitation assay with 2Flag-KAT14 and 2HA-NLS-Actin by using HA (upper panel) or Flag-beads (middle panel). AC40 antibody was used to blot endogenous actin in co-immunoprecipitation assay with or without overexpressed 2Flag-KAT14 by Flag-antibody (lower panel). IP, immunoprecipitation sample. Molecular weights indicated on the left. B. Western blots of co-immunoprecipitation with 2Flag-KAT14 and indicated HA-tagged actin mutants (upper panel). Lower panel shows the quantification of the relative amounts of actin mutants co-immunoprecipitated with KAT14. Data is normalized to wild type actin, and is the mean from 3 independent experiments with error bars showing standard deviation (S.D.). C. Western blots of co-immunoprecipitation assay with 2HA-actin and indicated Flag-KAT14 constructs. D. Western blots of pull down assay with actin, His-KAT14 and His-GST detected with the indicated antibodies. Note that for detecting the baits, unequal amounts were loaded on the SDS-PAGE gels (33 % of His-KAT14 sample and less than 1 % of His-GST) for visualization purposes. See supplementary figure 2 (S2C) for equal loading.

To study if actin binds directly to KAT14, we did a pull down with purified actin and recombinant His-tagged KAT14 expressed and purified from insect cells. Actin was detected on beads coated with KAT14, but not those coated with GST (Figure 5D), showing that the interaction between KAT14 and actin is direct. The Co-IP experiments with KAT14 truncations suggested that actin would interact with the C-terminus of KAT14. Since this part of KAT14 contains also the histone acetyl transferase (HAT) activity (Ma et al., 2017), it prompted us to investigate if actin-binding to KAT14 would affect its HAT activity. Using purified proteins, addition of KAT14 to core histones results in marked increase in acetylation of histone 4 lysine 5 (H4K5Ac) (Figure 6A,B), previously shown to be target for KAT14 (Guelman et al., 2009). Purified actin alone did not increase H4K5Ac above negative control (Figure 6A,B). However, when added together with KAT14, actin significantly decreased the acetylation efficiency of KAT14 (Figure 6A,B), demonstrating that *in vitro*, actin influences the KAT14 HAT activity. To study if actin could also regulate histone acetylation in cells, we overexpressed NLS-actin utilizing the inducible Strep-HA-NLS-actin cell line used for the AP-MS and measured H4K5Ac levels. In agreement with *in vitro* studies, overexpression of NLS-actin led to a significant decrease in H4K5Ac levels (Figure 6C,D). In line with our MS (Supplementary table 1) and co-IP experiments (Figure 5B), overexpression of NLS-R62D-actin lead to a even greater decrease in H4K5Ac levels (Figure 6C,D), further supporting the notion that KAT14 seems to prefer binding to monomeric actin.

**Figure 6.**
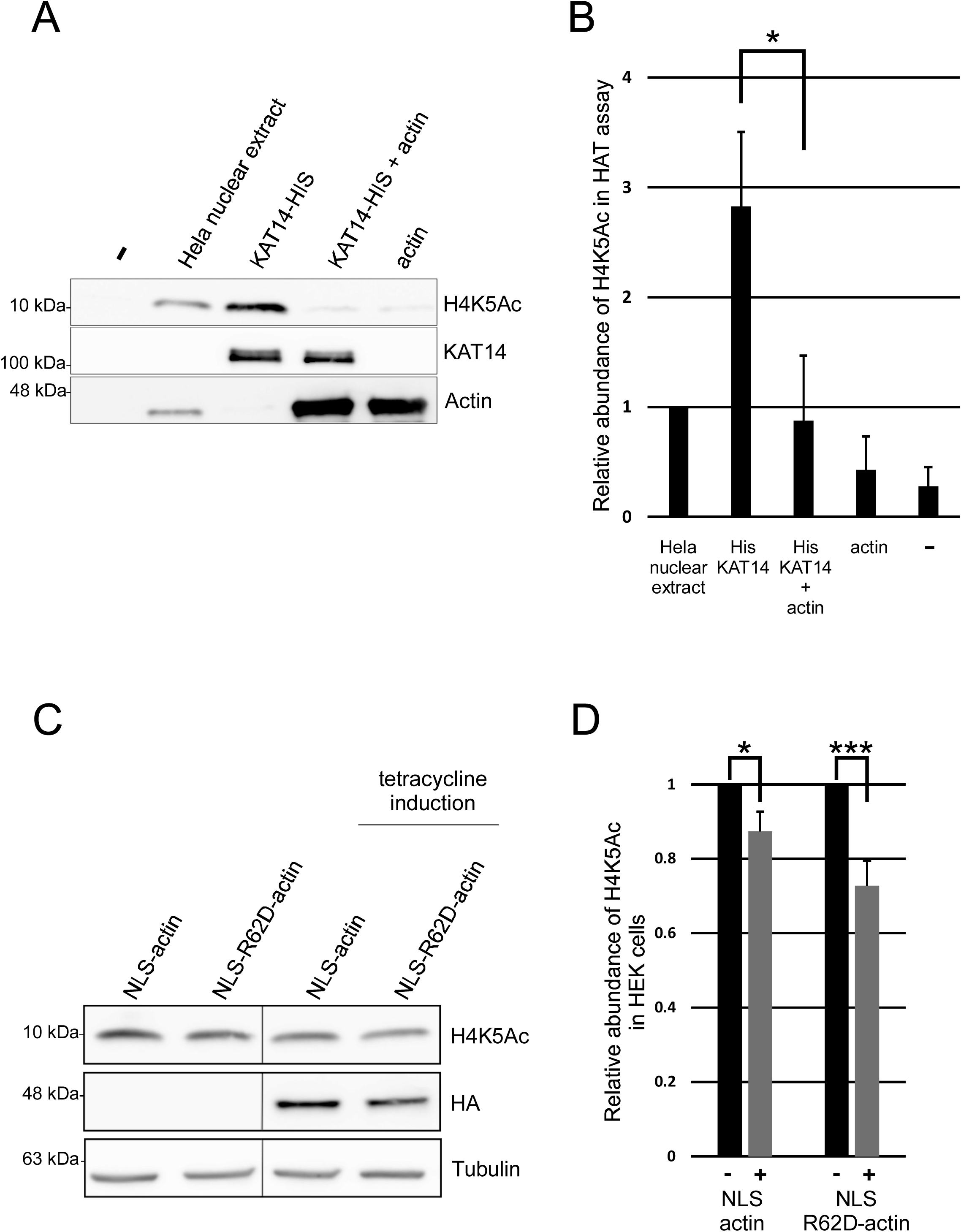
Actin inhibits KAT14 mediated H4K5 acetylation *in vitro* and in cells. A. Western blots of HAT assay with Hela nuclear extract, His-KAT14 without and with addition of actin and actin probed with the indicated antibodies. B. Quantification of the relative H4K5Ac abundance from western blots in 5A. Data is normalized to HeLa nuclear extract and mean from three independent experiments with error bars S.D. Statistics with two sided student’s t-test show significance between indicated samples (P=0.021). C. Western blots from total cell lysates of non-induced and tetracycline-induced stable cell lines expressing inducible HA-Strep-NLS-actin or HA-Strep-NLS-R62D-actin with the indicated antibodies. D. Quantification of the relative H4K5Ac abundance from western blots in 5C. Data is normalized to the non-induced sample, and is mean from three independent experiments with error bars S.D. Statistics with one sided student’s t-test show significance between indicated samples (respectively P=0.035 and P=0.008).

### Actin has a functional role in RNA splicing

Previous studies that have mainly focused on the hnRNP proteins (Obrdlik et al., 2008; Percipalle et al., 2002; Percipalle et al., 2001) have linked actin to pre-mRNA processing, and it has even been suggested that actin would follow the mRNA from the nucleus to the polysomes for translation [reviewed in (Percipalle, 2009)]. However, the functional role for actin in pre-mRNA processing, beyond the role in transcription, has not been demonstrated. Our BioID hits (Figure 3A,B) further strengthen the link between actin and pre-mRNA processing, but the fact that we recovered also core subunits of the spliceosome suggests that actin could play a role in regulating the splicing process itself. To examine this further, we first utilized the BiFC assay to confirm selected interactions between actin and splicing factors. All four tested splicing related factors SF3a120, SF3a60, SF3b49 and CDC5L gave fluorescent BiFC-signal with actin in the nucleus, (Figure 7A). Intriguingly, the BiFC-signal between CDC5L and actin showed a punctate appearance (Figure 7A), possibly suggesting that these proteins interact in a specific nuclear compartment. To test the functional significance of nuclear actin for mRNA splicing, we altered nuclear actin levels by importin 9 (Ipo9) (Dopie et al., 2012) and exportin 6 (Exp6) (Bohnsack et al., 2006; Stuven et al., 2003) depletion, thus decreasing and increasing nuclear actin levels, respectively, and performed survival of motor neuron 1/2 (SMN1/2) minigene splicing assay (Roca and Krainer, 2009). Interestingly, depletion of either Ipo9 or Exp6 increased exon 7 inclusion in the SMN2 minigene (Figure 7B). This indicates that alterations in nuclear actin dynamics disturb alternative exon skipping.

**Figure 7.**
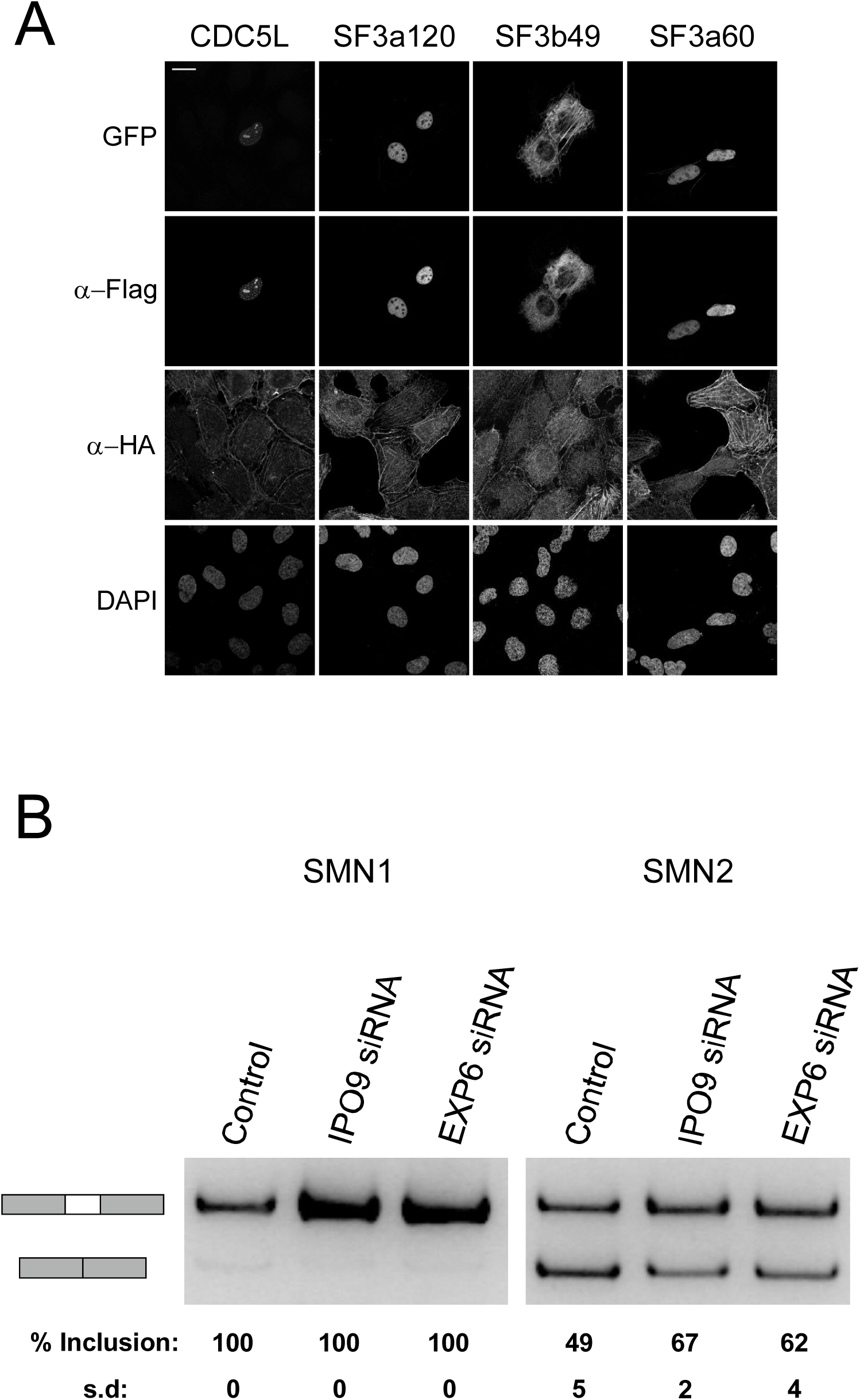
Actin associates with RNA splicing factors and disturbance of nuclear actin dynamics affect SMN2 alternative exon skipping. A. Confocal microscopy images of BiFC assay between different splicing factors and actin. Tetracycline induced HAGFP1-10-actin U2OS cells transiently transfected with the indicated FlagGFP11 constructs. Data shown as in Figure 4B. B. RT-PCR analysis of SMN1/SMN2 minigene splicing from control, Ipo9, and Exp6 siRNA treated NIH 3T3 cells. The SMN1 and SMN2 mRNA products are indicated on the left. Quantification from three independent experiments as mean percentage of exon inclusion with S.D indicated below. Difference between control siRNA and Ipo9 siRNA exon 7 inclusion is significant (P=0.03).

## Discussion

Despite its essential functions in the cell nucleus, we still understand relatively poorly how actin operates in this cellular compartment at the molecular level. In the cytoplasm, actin has hundreds of binding partners, and many of these interactions have been characterized in molecular, and even structural, detail. As a step towards understanding the mechanisms by which actin operates in the nucleus, we have here aimed to identify its nuclear binding partners by using two complimentary MS-methods, AP-MS and BioID. Our analysis identifies already established binding partners for actin, such as components of chromatin remodeling complexes, and provides new insights into known actin-regulated processes, such as transcription. In addition, we identify completely novel binding partners and functions for actin in the nucleus.

The abundant cytoplasmic actin pool, as well as biochemical properties of actin itself, pose significant challenges for identifying nucleus-specific interactions for this protein. Cell fractionation is a common approach to study compartment-specific properties of proteins. Based on our own experiences, actin is quite hard to fractionate reliably, and the abundant cytosolic actin is easily contaminating the much less-abundant nuclear fraction. In addition, actin polymerization under physiological salt conditions further complicates the issue. Here we applied two approaches to overcome these issues: 1) increasing the actin amounts in the nucleus by fusing actin with an NLS and 2) comparing the interactomes to actin without the NLS. Indeed, GO term analysis revealed that our approach enriched proteins with the GO term nucleus (GO: 0005634; Figure 1B), but our approach has also its limitations. One limitation is that we are utilizing tagged actin for both AP-MS and BioID. In the case of AP-MS the tag is equivalent to an epitope tag (less than 4 kDa), but the mutated biotin ligase (BirA*) used in BioID is approximately 39 kDa. It is a valid concern that the tag may prevent some interactions of actin, and thus render them undetectable with the approach taken here. For example, although components of the SWI/SNF and SRCAP/TIP60 chromatin remodeling complexes were readily detected in our nuclear actin interactomes (Figure 2B,C), no specific subunits of the hINO80 complex scored as high-confidence interactions in our MS screens (Supplementary table 2). Earlier studies with yeast INO80 (Kapoor et al., 2013; Shen et al., 2003) and *in vitro* biochemical studies with actin, Arp4 and Arp8 (Fenn et al., 2011) clearly establish the presence of actin in this complex. Recent structural work of the Arp8 module, containing also actin and Arp4 (Knoll et al., 2018) may explain this discrepancy. In INO80, actin is sandwiched between Arp4 and Arp8, while in SWI/SNF and SRCAP/Tip60 complexes, actin pairs only with Arp4. Tagging actin could therefore either directly inhibit binding to the INO80 or alternatively, actin integrates more stably to the INO80 complex than to the other actin-containing remodelers, and the tagged-actin cannot compete the endogenous actin out of the complex. Development of methods that allow studies on the endogenous nuclear actin interactions are therefore needed in the future to overcome these limitations. Another limitation of our experimental set up relates to the use of the wild type actin without the NLS to eliminate the cytoplasmic interactions from the nuclear interactomes. As we have shown before, actin shuttles in and out of the nucleus (Dopie et al., 2012), and hence the wild type actin also displays some nuclear localization (Supplementary figure 1), and consequently will interact with nuclear proteins to a certain extent. In line with this, especially the high-confidence interactions obtained with AP-MS for actin contained a substantial portion (38%) of hits with the GO term nucleus (Figure 1B). This tendency was most notable with the non-polymerizable actin mutant R62D-actin (Supplementary table 1), and can be explained by the molecular properties of R62D-actin: first it cannot be incorporated into the actin cytoskeleton as tightly as WT actin (Posern et al., 2002) and second, it has higher nuclear import rate than WT actin (Dopie et al., 2012). Hence the analysis against the wild type actin may eliminate especially those interactions that take place both in the cytoplasm and in the nucleus. This could explain the relatively low number of known actin-binding proteins that were detected as high-confidence interactors for nuclear actin (Figure 2C). Recent studies have finally shown that nuclear actin can polymerize into canonical actin filaments (Grosse and Vartiainen, 2013), raising the question on how polymerization influences nuclear actin interactions. Although answering this question was not an aim of this study, we utilized the non-polymerizable actin mutant R62D-actin alongside wild type actin in our MS screen. However, we did not observe major differences between their nuclear interactomes (Figure 1C). Indeed, most of the known actin interaction partners in the nucleus, including Mrtf-A (Vartiainen et al., 2007), pTEF-β (Qi et al., 2011), and hnRNPU (Kukalev et al., 2005), as well as the different chromatin remodeling complexes (Kapoor et al., 2013; Zhao et al., 1998) seem to bind monomeric actin. It must also be noted that our experimental set up was not geared towards identifying binding partners for nuclear filamentous actin. It is unlikely that the filaments would stay intact during the AP-MS procedure, and the long biotinylation time required in BioID would not be compatible with the transient nature of the nuclear actin filaments (Baarlink et al., 2017; Baarlink et al., 2013; Plessner et al., 2015). Nevertheless, this is an important avenue for future studies, since nuclear actin polymerization seems to have a key role both during cell cycle (Baarlink et al., 2017) and in DNA damage response (Caridi et al., 2018; Schrank et al., 2018).

Beyond its roles in chromatin remodeling complexes, actin has also been linked to different steps of transcription (Fomproix and Percipalle, 2004; Hofmann et al., 2004; Kukalev et al., 2005; Obrdlik et al., 2008; Percipalle et al., 2003), and our recent genome-wide analysis indicates that actin interacts with basically all transcribed genes (Sokolova et al., 2018). In support of this, multiple proteins related to transcription from the assembly of the PIC to Pol II-mediated elongation (Jonkers and Lis, 2015) were identified as high-confidence interactions in our MS screens (Figure 3A,B, Supplementary table 2). Notably, almost all of these transcription-related hits were obtained with BioID (Figure 3A,B, Supplementary table 2), suggesting that these interactions may be rather transient and dynamic. Alternatively, they could be very context-specific, and thus not retained in the AP-MS. Unexpectedly, we did not obtain a single subunit of the core RNA polymerases as a high-confidence interactor in our assays (Supplementary table 2), despite the fact actin was shown to co-purify with Pol II already decades ago (Egly et al., 1984). Whether this is due to the experimental set up, for example the usage of tagged-actin, awaits further studies. Beyond gene expression, recent studies have linked actin to also maintenance of genomic integrity via movement of damaged DNA (Caridi et al., 2018; Schrank et al., 2018) and DNA replication (Parisis et al., 2017). Even though our mass spec analysis was done in normal, cycling cells, few of the putative interaction partners for actin were related to DNA replication and DNA damage response (Figure 3A, Supplementary table 2). In the future, this nuclear actin interactome analysis could be expanded to different cell cycle phases and to samples with induced DNA damage.

It is well established that actin associates with nuclear Arps to bind chromatin remodelers and modifiers like SWI/SNF, SRCAP/Tip60 and INO80 [reviewed in (Kapoor and Shen, 2014)]. To our surprise, the shared hits from the AP-MS and BioID screens included several subunits from the hATAC, a histone modifying complex without nuclear Arps (Guelman et al., 2009), and our further analysis revealed direct interaction between the histone acetyl transferase KAT14 and actin (Figure 5D). We found that actin interacts with the C-terminus of KAT14, which also contains the HAT activity, and actin inhibited the KAT14 HAT activity *in vitro* (Figure 5C, 6A,B). The hATAC contains also another HAT, GCN5, and it seems that the HAT activities of both GCN5 and KAT14 contribute to the total HAT activity of the ATAC complex (Guelman et al., 2009; Suganuma et al., 2008). However, KAT14 and GCN5 seem to have distinct histone substrate preferences (Suganuma et al., 2008). In the future, it will be interesting to study if actin regulates the activity of also the whole hATAC complex, or whether its functions are restricted to KAT14. We were also able to detect significant decrease in H4K5 acetylation levels, which have previously been shown to be dependent on KAT14 (Guelman et al., 2009), in cells overexpressing NLS-actin or even more prominently in cells overexpressing NLS-R62D actin (Figure 6C,D). However, we cannot exclude the possibility that these cellular effects are due to actin’s actions on some other proteins that control histone acetylation. Indeed, a recent paper demonstrated that actin monomers interact with histone deacetylases (HDAC) 1 and 2, and that polymerizing nuclear actin increased HDAC activity and decreased histone acetylation. However, the interaction between actin and HDAC proteins did not seem to be direct (Serebryannyy et al., 2016a). Further investigations are needed to reveal how actin is involved in controlling the delicate histone acetylation balance in cells. Interestingly, KAT14 has also been demonstrated to promote smooth muscle gene expression by interacting with SRF, CRP2 and myocardin (Ma et al., 2017). It would be interesting to investigate whether KAT14 also plays a role in controlling the actin-regulated myocardin family member, MRTF-A (Vartiainen et al., 2007).

Intriguingly, we obtained numerous RNA splicing related hits with our nuclear actin BioID screen (Figure 3A,B) and these contained multiple subunits from the major spliceosome complexes (Wahl and Luhrmann, 2015). In the BiFC assay, one of the tested splicing factors, CDC5L, displayed a punctate interaction pattern with actin in the nucleus (Figure 7A), which might indicate that the interaction occurs in nuclear speckles, domains known to be enriched in pre-mRNA splicing factors. Since splicing occurs co-transcriptionally (Herzel et al., 2017), the splicing related hits could be explained by the association of actin with the transcription machinery (Figure 3B). However, we were able to show that alterations in the nuclear actin levels disturb alternative splicing of the SMN2 minigene (Figure 7B), potentially suggesting a functional role for actin in this process. However, we must note that since alterations in nuclear actin levels are also linked to decreased transcription (Dopie et al., 2012), this effect could be indirect, e.g. actin could be required for transcription of a spliceosomal component. Further experiments, such as deep RNA sequencing, are required to elucidate the exact role of actin in transcription vs. RNA processing. We have earlier shown that depletion of certain splicing related factors, such as CDC73, results in the formation of nuclear actin filaments, or bars (Rohn et al., 2011). It is therefore tempting to speculate that actin and RNA splicing are connected and that they both influence each other.

Taken together, our interactome analysis opens new avenues for studying the mechanisms by which actin operates in the nucleus by providing both new angles for known actin-regulated nuclear processes and completely novel interactions as well as functions. The next step will be to analyze these interactions preferably in biochemical detail. This may prove challenging, since many interactions of actin, for example in the context of transcription, may be highly dynamic and transient, or could be highly context-specific, making it difficult to reconstitute these *in vitro.* However, biochemical data for nuclear actin interactions is crucial, so that we can start to manipulate the interactions and functions separately. This is critical, because this interactome analysis has further highlighted the multifunctional nature of nuclear actin.

## Materials and methods

### Plasmids

Detailed information of the cloned plasmids are available upon request. Majority of the plasmids generated here will be available at Addgene and Addgene ID number is indicated after the plasmid.

To generate AP-MS compatible Strep-HA-actin constructs we used Gateway (GW) compatible N-terminally tagged destination vector pcDNA5/FRT/TO/SH/GW (Glatter et al., 2009) and actin constructs [pENTR-actin (118378), pENTR-NLS-actin (118380), pENTR-R62D-actin (118379), pENTR-NLS-R62D-actin (118381)]. To generate BioID compatible N-terminally tagged HA-BirA* destination vector (HA-BirA*-pDEST-N-pcDNA5/FRT/TO, 118375), pcDNA5/FRT/TO/SH/GW was altered by replacing the Strep-tag with BirA* from pcDNA 3.1 Myc-BirA (Roux et al., 2012) with restriction digestion. GW compatible vectors used in Bimolecular Fluorescence Complementation (BiFC) assay [HA-GFP1-10-actin-pcDNA4/TO (118370) and Flag-GFP11-pDEST] were cloned from pmGFP1-10 and pmGFP11 plasmids (Brunello et al., 2016), which were kind a gift from Henri Huttunen. GW cloning compatible destination vectors containing N- or C-terminal tags [2HA-pDEST-N/C (118373, 118374), 2Flag-pDEST-N/C (118371, 118372), Flag-GFP11-pDEST-N/C (118366, 118367), HA-GFP1-10-pDEST-C/N (118368, 118369)] were generated by restriction digestion cloning to pDEST27 vector (Thermo Fisher Scientific) as backbone. AP-MS or BioID hits were cloned to Flag-GFP11-pDEST by using GW cloning and entry vectors from Human ORFeome collection, except BRG1/SMARCA5, which was cloned from Hela cell cDNA. KAT14 was also cloned with GW to 2Flag-pDEST and to pDEST10 (KAT14-pDEST10, 118382) (Thermo Fisher Scientific). KAT14 truncations [pENTR-KAT14-1-546 (118377), pENTR-KAT14-547-782 (118376)] were cloned to 2Flag-pDEST vector with GW cloning. PENTR-G168D,Y169D-actin (118384) and pENTR-V159N-actin (118383) plasmids were generated from the pENTR-actin plasmid by site mutagenesis and cloned to 2HA-pDEST vector with GW cloning. SMN1 and SMN2 minigene plasmids were kind gift from Mikko Frilander.

### Antibodies

The following antibodies were from Merck: Flag (F1804) (1:400), HA (H3663) (1:400), Tubulin (B512) (1:2500), Actin (A3853) (1:2000). The following antibodies were from Abcam: KAT14 (ab127040) (1:1000) and H4K5Ac (ab51997) (1:10 000). The following antibodies were from Santa Cruz: HA-Probe (sc-805) (1:300). The following secondary antibodies were from Thermo Fisher Scientific: HRP-conjugated anti-mouse (G-21040) (1:7500), HRP-conjugated anti-rabbit (G-21234) (1:7500), Alexa-Fluor-488-conjugated streptavidin (S11223) (1:500), Alexa-Fluor-594-conjugated anti-mouse (A11005) (1:500), Alexa-Fluor-594-conjugated anti-rabbit (A11012) (1:500), Alexa-Fluor-647-conjugated anti-mouse (A-31571) (1:500). The following secondary antibodies were from Merck: HRP-conjugated Flag M2 (A8592) (1:7500), HRP-conjugated HA (H6533) (1:7500), HRP-conjugated HIS (A7058) (1:7500).

### Cell lines and transfections

All cell lines (Hela, Flp-In™ T-REx™ 293 and U2OS) were cultured in DMEM (Merck) supplemented with 10% Fetal Bovine Serum (FBS) (Thermo Fisher Scientific), Penicillin-Streptomycin (Thermo Fisher Scientific) and maintained at 37°C and 5% CO_2_.

For generation of the stable cell lines with inducible expression of the BirA*- and Strep-HA-tagged versions of the different actin constructs (actin, R62D-actin, NLS-actin and NLS-R62D-actin), Flp-In™ T-REx™ 293 cell lines (Invitrogen, Life Technologies, R78007, cultured according manufacturer’s recommended conditions) were co-transfected with the expression vector and the pOG44 vector (Invitrogen) using the Fugene6 transfection reagent (Roche Applied Science). Two days after transfection, cells were selected in 15 μg/ml blasticidin S (Invivogen) and 50 μg/ml hygromycin (Thermo Fisher Scientific) for 2 weeks and emerged positive clones were tested and clones with correct actin localization and strong expression levels were amplified and used for experiments. Stable cells expressing BirA*- or Strep-HA-tag fused to green fluorescent protein (GFP) were used as negative controls and processed in parallel to the bait proteins.

Stable cell line used in BiFC assay was generated by transfecting HA-GFP1-10-actin-pcDNA4/TO plasmid to human osteosarcoma (U2OS) cells stably expressing tetracycline repressor by using lipofectamine 2000 (Thermo Fisher Scientific) with manufacturer’s protocol. Cells were selected with 300 μg/ml zeocin (Invivogen) and 2,5 μg/ml blasticidin S and the clones were screened for strong induction level and low background signal. For the BiFC assay, HA-GFP-1-10-actin U20S cells were plated onto 24-well tissue culture plate at a density of 12,000 cells per well. The next day, cells were induced with 1:1000 dilution of tetracycline. On day three cells were transfected with DNA constructs using JetPrime transfection reagent (Polyplus transfection), according to the manufacturer’s instructions. In total, 250 ng of DNA was used for every transfection reaction, with the experimental vector amount 25 ng. The pEF-vector was used to fill up the total amount of DNA.

### Immunoprecipitations

For immunoprecipitation (IP), Hela cells were plated onto 10 cm culture dish at of 1,000,000 cells per dish. The next day, cells were transfected with 2Flag-KAT14, 2Flag-KAT14 truncations and 2HA-actin constructs (R62D-actin, V159N-actin and G168D,Y169D-actin) using JetPrime transfection reagent (Polyplus transfection), according to the manufacturer’s instructions. Equal amount (3 μg) of expression vectors was used for transfection. 48 h after transfection cells were collected and lysed in lysis buffer (50 mM Tris-HCl, pH 8, 100 mM NaCl, 1 % IGEPAL and protease inhibitors (Roche)). Cleared lysate was obtained by centrifugation and diluted 1:2 in IP buffer [50 mM Tris-HCl, pH 8, 100 mM NaCl, and protease inhibitors (Roche)]. Prewashed 30 μl anti-flag^®^ M2 affinity gel (Merck) was added to diluted lysates and after 3 h on rotation at +4 °C, the samples were washed three times with IP-buffer, and bound proteins eluted with 90 μL of 1 × SDS-PAGE loading buffer. Samples (33% of the immunoprecipitates and 10% of the inputs) were separated in 10% SDS-PAGE, transferred onto nitrocellulose membrane and the proteins detected with directly conjugated anti Flag M2-peroxidase, anti-HA HA-7 peroxidase, or with mouse anti-Actin.

### MS sample preparation and MS analysis

Strep-HA or HA-BirA* stable cell lines were expanded to 80% confluence in 6 × 150 mm cell culture plates. For the AP-MS approach, 1 μg/ml tetracycline was added for 24 h induction of protein expression before harvesting the cells. For BioID, in addition to tetracycline, 50 μM biotin (Thermo Fisher Scientific) was added 24 h before harvesting. Cells from confluent plates (~6 × 107 cells) were pelleted as one biological sample, and at least two biological replicates were prepared for each bait. Cell pellets were snap frozen and stored at −80°C until sample preparation.

For AP-MS, the cell pellet was lysed in 3 ml of lysis buffer 1 (1% IGEPAL, 50 mM Hepes, pH 8.0, 150 mM NaCl, 50 mM NaF, 5 mM EDTA, supplemented with 0.5 mM PMSF and protease inhibitors; Roche). For BioID, the cell pellet was lysed in 3 ml lysis buffer 2 (1 % IGEPAL, 50 mM Hepes, pH 8.0, 150 mM NaCl, 50 mM NaF, 5 mM EDTA, 0.1% SDS, supplemented with 0.5 mM PMSF and protease inhibitors; Roche). Lysates were sonicated and treated with benzonase (Merck). Cleared lysate was obtained by centrifugation and loaded consecutively on spin columns (Bio-Rad) containing lysis buffer 1 prewashed 200 μl Strep-Tactin beads (IBA, GmbH). The beads were then washed 3 ×1 ml with lysis buffer 1 and 4 × 1 ml with wash buffer (50 mM Tris-HCl, pH 8.0, 150 mM NaCl, 50 mM NaF, 5 mM EDTA). Following the final wash, beads were then resuspended in 2 × 300 μl elution buffer (50 mM Tris-HCl, pH 8.0, 150 mM NaCl, 50 mM NaF, 5 mM EDTA, 1 mM Biotin) for 5 min and eluates collected into an Eppendorf tubes. Samples were snap frozen and stored at −20°C until MS-LC analysis.

MS-LC samples were treated with 5 mM Tris (2-carboxyethyl)phosphine (TCEP) and 10 mM iodoacetamide for 30 mins at 37 °C to reduce cysteine bonds and alkylation, respectively. The proteins were then digested to peptides with sequencing grade modified trypsin (Promega, V5113) at 37 °C overnight. After quenching with 10% TFA, the samples were desalted by C18 reversed-phase spin columns according to the manufacturer’s instructions (Harvard Apparatus). The eluted peptide sample was dried in vacuum centrifuge and reconstituted to a final volume of 30 μl in 0.1% TFA and 1% CH3CN.

The MS analysis was performed on Orbitrap Elite hybrid mass spectrometer coupled to EASY-nLC II-system using the Xcalibur version 2.7.0 SP1 (Thermo Fisher Scientific). 4 μl of the tryptic peptide mixture was loaded into a C18-packed pre-column (EASY-Column™ 2 cm × 100 μm, 5 μm, 120 Å, Thermo Fisher Scientific) in 10 μl volume of buffer A and then to C18-packed analytical column (EASY-Column™ 10 cm × 75 μm, 3 μm, 120 Å, Thermo Fisher Scientific). Sixty-minute linear gradient at the constant flow rate of 300 nl/minute from 5 to 35% of buffer B (98% acetonitrile and 0.1% formic acid in MS grade water) was used to separate the peptides. Analysis was performed in data-dependent acquisition: one high resolution (60,000) FTMS full scan (m/z 300–1700) was followed by top20 CID-MS2 scans in ion trap (energy 35). Maximum FTMS fill time was 200 ms (Full AGC target 1,000,000) and the maximum fill time for the ion trap was 200 ms (MSn AGC target of 50,000). Precursor ions with more than 500 ion counts were allowed for MSn. To enable the high resolution in FTMS scan preview mode was used.

### Search Parameters and Acceptance Criteria

Proteins were identified using Proteome Discoverer™ software with SEQUEST search engine (version 1.4, Thermo Fisher Scientific). Thermo .raw files were searched against the human component of the UniProt-database (release 2014_11; 20130 entries) complemented with trypsin, BSA, GFP and the tag sequences. Trypsin was used as the enzyme specificity. Search parameters specified a precursor ion tolerance of 15 ppm and fragment ion tolerance of 0.8 Da, with up to two missed cleavages allowed for trypsin. Carbamidomethylation (+57.021464 Da) of cysteine residues was used as static modification whereas oxidation (+15.994491 Da) of methionine and biotinylation (+226.078 Da) of lysine residues or N terminus were used as dynamic modification. Peptide false discovery rate (FDR) was calculated using Percolator node of software and set to <0.01. Spectral counting was used to produce semiquantitative data. Identification metrics for each sample are listed in Supplementary table 1.

### Identification of High-confidence interactions

Significance Analysis of INTeractome (SAINT)-express version 3.6.0 (Teo et al., 2014) and Contaminant Repository for Affinity Purification (CRAPome, http://www.crapome.org/) (Mellacheruvu et al., 2013) were used for identification of specific high-confidence interactions (HCI) from our MS data. 11 GFP-Strep-HA runs [one GFP-Strep-HA control and 10 CRAPome GFP-Strep-HA controls (CC295, CC306, CC307, CC411, CC316, CC323, CC328, CC329, CC330, CC332)] were used as control counts for each AP-MS hit. 4 HA-BirA*-GFP runs (from 2 biological HA-BirA*-GFP replicates) were used as control counts for each BioID-MS hit. Actin-MS and R62D-actin MS replicates (total 4 in AP-MS and in BioID-MS) were combined to analyze the AP-MS and BioID actin interactomes. NLS-actin-MS and NLS-R62D-actin MS replicates (total 4 in AP-MS and total 6 in BioID-MS) were combined to analyze nuclear actin AP-MS and BioID interactomes. Identified proteins were considered as high confidence interactions if they were identified in at least two biological replicates, and had average spectral count fold change ≥ 2 (in AP-MS) or ≥ 0.5 (in BioID) compared to control. In addition, the high confidence interactions for nuclear actin had average spectral count fold change ≥ 1 (in AP-MS) or ≥ 0.5 (in BioID) compared to the respective actin sample. For SAINT-express analysis spectral counts of different hit candidates (2 or 3 biological replicates) were pooled and analyzed over control with SAINT Express. The final results only consider proteins with SAINT score ≥ 0.88. This corresponds to an estimated protein-level Bayesian FDR of <0.01. Endogenously biotinylated proteins such as Acetyl-CoA carboxylase 2 (ACACB), Acetyl-CoA carboxylase 1, (ACACA) as well as proteins from other species than homo sapiens were removed from the final HCI list. Database for Annotation, Visualization and Integrated Discovery (DAVID) annotation tool (Huang et al., 2009) was used to create GO term tables from the hit lists. Shared hits and specific AP-MS hits were show with STRING map (version 10.5) (Szklarczyk et al., 2017).

### Microscopy

Cells on coverslips were fixed with 4% paraformaldehyde for 15 min, washed three times with PBS and permeabilized for 7 min with 0,2% TritonX-100 (Merck) in PBS. For antibody staining, permeabilized cells were blocked with blocking buffer (1% gelatin (Merck), 1 % BSA (Merck) and 10 % FBS (Lonza) in 1 × PBS) for 1 h and incubated with primary antibody for 1 h. Coverslips were washed and incubated with Alexa Fluor conjugated secondary antibody for 1h. Coverslips were washed three times with 1 × PBS, once with MQ water and mounted in Prolong Diamond with DAPI (Thermo Fisher Scientific).

Wide-field fluoresce microscope Leica DM6000 (Leica, Welzlar, Germany) with HCXPL APO 63x/1.40–0.60 oil objective and confocal microscope Leica TCS SP8 STED 3X CW 3D with HC PL APO 93x/1.30 glycerol objective were used to image the samples. The image files were processed with Leica LAS X and ImageJ software.

### Insect cell expression and protein purification

Recombinant human His-KAT14 was produced by using the MultiBac baculovirus expression vector system. Briefly, KAT14-pDEST10 plasmid was transformed into DH10MultiBac cells and recombinant bacmids were isolated and the presence of the coding sequence of KAT14 was confirmed with PCR. A bacmid containing the right sequence was transfected into Spodoptera frugiperda cells (Sf9) by using Fugene HD (Promega). The baculoviruses were amplified and used for protein expression. The expression of recombinant human KAT14 was conducted by infection 2 × 106 Sf9 cells with the baculoviral stock at MOI 1. The cells were harvested after 72 h and the cell pellets were snap frozen and stored at −80 °C

Cell pellets were resuspended in lysis buffer (25 mM Tris-HCl pH 8.0, 10 mM NaCl, 1mM MgCl2, 2 mM B-mercabtoethanol, 0,2 % IGEPAL, 1 mM PMSF, protease inhibitors (Roche)) with Benzonase (Merck) and lysed with EmulsiFlex-C3 (AVESTIN) with gauge pressure approximately 15000 psi for 10 min. Immediately after lysing NaCl concentration was adjusted to 150 mM and also 10 mM Imidazole was added to cells extracts. Cell extract was incubated for 10 min at +4 °C and then clarified by spinning at 92000 × g for 30 min and immediately processed to metal (Ni2+) affinity purification. The cell extract from about 1 liter of infected cells was mixed with Ni2+-NTA agarose beads (Qiagen) that were equilibrated with lysis buffer. Beads were incubated with the extract for 2 h in rotation at +4 °C. The beads were washed three times with the buffer A (25 mM Tris-HCl pH 8.0, 300 mM NaCl, 2 mM mM B-mercaptoethanol) and two times with buffer A with addition of 10 mM Imidazole. The last wash was conducted in a disposable 10 ml Poly-prep column (Bio-RAD) and the protein was eluted from the beads with 500 mM Imidazole in lysis buffer. The protein was further purified with gel filtration column Superdex 200 (Superdex 200 HiLoad 16/60, Pharmacia), which was equilibrated with (25 mM Tris-HCl pH 7.5, 150 mM NaCl, 0,5 mM EDTA, 2 mM B-mercaptoethanol, 10 % glycerol). Fractions were analyzed by SDS-PAGE, and fractions containing His-KAT14 were concentrated and stored at −80 °C.

### Pulldown with purified proteins

His-GST lysate (500 μl) and purified His-KAT14 were immobilized on Ni2+-NTA agarose beads equilibrated with storage buffer (25 mM Tris-HCl pH 7.5, 150 mM NaCl, 0,5 mM EDTA, 2 mM B-mercaptoethanol, 10 % glycerol) (30 μl, Qiagen) for 2 hours on rotation at + 4 °C. Beads were washed once with Binding buffer 1 (25 mM Tris-HCl pH 7.5, 150 mM NaCl, 1 mM DTT), three times with Binding buffer 2 (25 mM Tris-HCl pH 7.5, 500 mM NaCl, 1 mM DTT) and still once with G-Buffer (5 mM Tris-HCl pH 7.5, 0,2 mM ATP, 0,2 mM CaCl2, 0,5 mM DTT). After washes beads were incubated with 4-fold amount of rabbit skeletal muscle actin compared to HIS-KAT14 in G-buffer for 3 hours on rotation at + 4 C. Rabbit skeletal muscle actin was prepared as described in (Pardee and Spudich, 1982). After incubation the beads were washed three times with G-buffer and bound proteins were eluted with 1 × SDS-PAGE loading buffer. Samples were boiled for 5 min and the proteins were separated in 7,5 % Mini-PROTEAN^®^ TGX™ Precast Protein Gel (BioRAD) and electroblotted to nitrocellulose membrane. The membrane was probed with KAT14, Actin, or His antibodies to assess their association with His-fusions.

### HAT assays in vitro and in cells

Core histones (5 μg; Merck) were incubated with 30 μM Acetyl CoA in different conditions: with MQ water, with 20 μg Hela nuclear extract, with 250 nM His-KAT14, with 250 nM His-KAT14 with addition of 2,5 μM actin and with 2,5 μM actin alone. 25 μl reactions were performed in 1 × HAT buffer (50 mM Tris-HCl, pH 8.0, 50 mM NaCl, 0.1 mM EDTA, 0.01% Igepal) and incubated in +30 °C for 1 h. After incubation the proteins were denaturated in 1 × SDS loading buffer and boiled for 5 min before separation by 7,5 % Mini-PROTEAN^®^ TGX™ Precast Protein Gel and electroblotted to nitrocellulose membrane or stained with Coomassie. The membrane was probed with KAT14, Actin and H4K5Ac antibodies. H4K5Ac was normalized to H4 protein amount and to Hela nuclear extract ratio, which was set to correspond 1. Error bars show the S.D. of 3 independent experiments. Significance between indicated samples shown with asterisk (*, P<0.05)

Strep-HA-NLS-actin and Strep-HA-NLS-R62D-actin cells were plated in 6 well plate wells in 250,000 cells per well. The next day half of the cells were induced with 1:1000 dilution of tetracycline. On day three, the all the cells were harvested and washed once with 1 × PBS before lysing into 1 × SDS loading buffer. Samples were boiled for 5 min and the proteins were separated in 7,5 % Mini-PROTEAN^®^ TGX™ Precast Protein Gel and electroblotted to nitrocellulose membrane. The membrane was probed with KAT14, Actin, Tubulin and H4K5Ac antibodies. Quantification of the ratio of relative H4K5Ac amount was normalized to tubulin and to non-induced NLS-actin or NLS-R62D-actin ratio, which was set to correspond 1. Error bars show the S.D. of 4 independent experiments. Significance between indicated samples shown with asterisk (*, P<0.05, ***, P<0.01)

### SMN1/SMN2 minigene splicing assay

For splicing assay NIH cells were plated onto 6-well tissue culture plate at a density of 40,000 cells per well. The following day, cells were transfected with 20 nmol siRNA (Ipo9: 5’-CACCGAGGAGCAGAUUAAA-3’, Exp6 5’-CGUUGAUAUUGGACGCCAA-3’ from Sigma; negative control: AllStars negative control from Qiagen) using Interferin transfection reagent (Polyplus transfection), according to manufacturer’s instructions. On day 4, cells were re-transfected with siRNAs and with SMN1 and SMN2 minigene plasmids (75 ng of SMN minigene, total DNA amount 1 μg per well added up with mock DNA) [previously described in (Roca and Krainer, 2009)] with JetPrime transfection reagent (PolyPlus transfection). The following day total RNA was extracted with Nucleospin RNA II kit instructed by the manufacturer (Macherey-Nagel). Total RNA (500 ng) was used for complementary DNA synthesis using Maxima First Strand cDNA Synthesis Kit (Thermo Fisher Scientific). cDNAs from the SMN1/2 minigenes was amplified with pCI-Fwb and pCI-Rev primers (Cartegni et al., 2006) with Phusion Hot Start (Thermo Fisher Scientific), 25 cycles. PCR products were separated with 6% native PAGE stained with 0,5 μg/ml EtBr. Gel images were acquired with Gel Doc™ EZ System (BioRAD) and the intensity of the bands corresponding to different splice variants were measured. In all cases, the standard deviations were ≤ 5%, such that the exon-inclusion percentage values can be compared between experiments. Difference between control siRNA and Ipo9 siRNA exon 7 inclusion is significant (P <0.05).

### Statistical Analyses

Statistical analyses were performed in Excel. The data for the HAT assays from the western blots and the SMN1/SMN2 splicing assay from native PAGE were analyzed by two-tailed student’s t-test because the data conformed to normal distribution. The data for Strep-HA-NLS-actin and Strep-HA-R62D-actin cell induction assays was analyzed with one sided student’s t-test. Significance was determined by P < 0.05, and the actual P values are indicated in the figure legends.

## Acknowledgements

We thank Paula Maanselkä and Mikko Honkanen for technical assistance, as well as Henri Huttunen and Mikko Frilander for plasmids. Mikko Frilander is also acknowledged for his help with the SMN minigene assays. Imaging was carried out at the Light Microscopy Unit (LMU) of the Institute of Biotechnology. This work was supported by the Academy of Finland, ERC, Sigrid Juselius and Jane & Aatos Erkko foundation grants to MKV.

---

**Supplementary figure 1 (S1).**
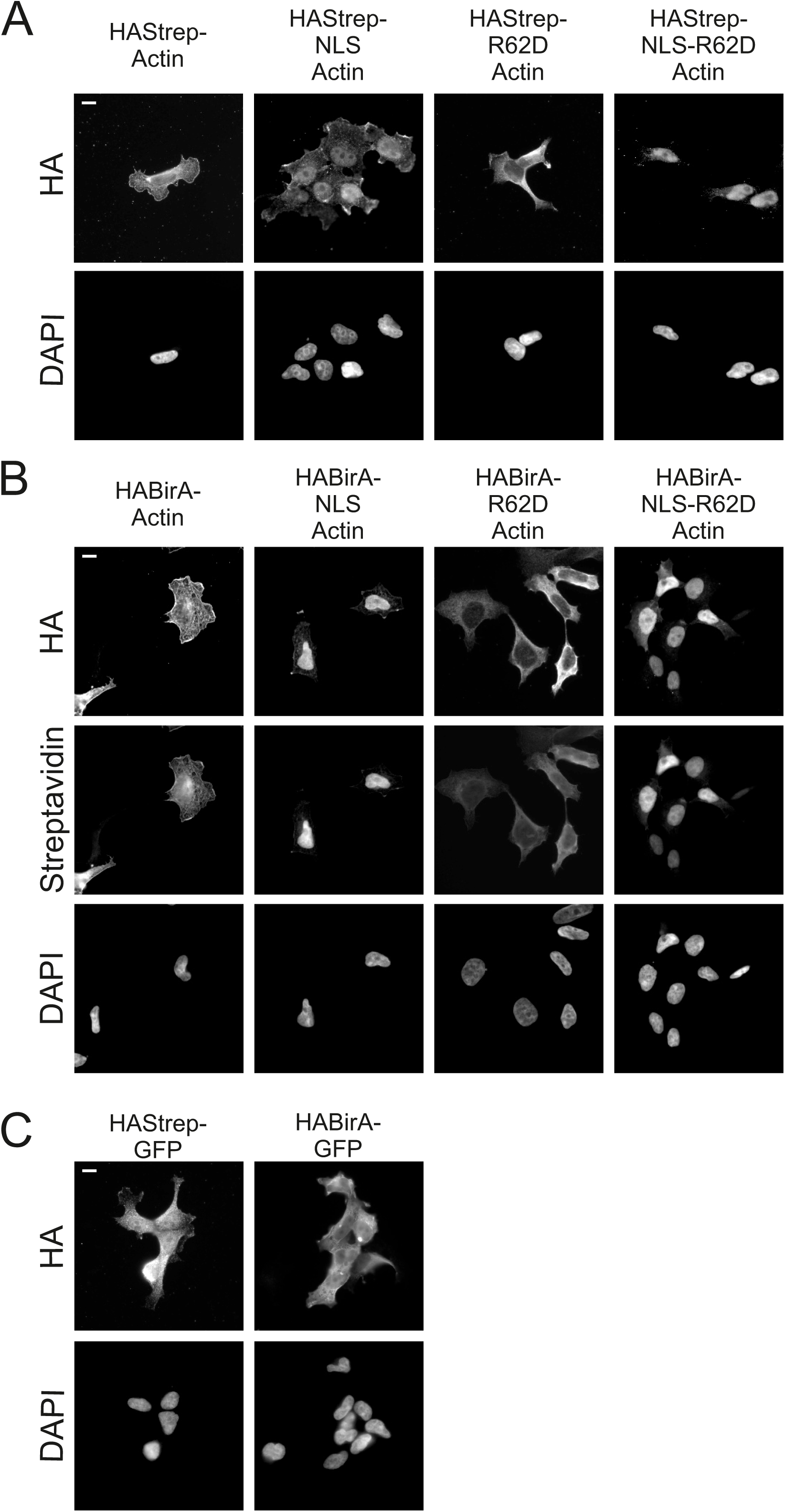
Stable cell lines used for the MS experiments. A. Wide-field microscopy images of stable, inducible cell lines expressing HAStrep-tagged actin constructs as indicated, and stained with HA antibody and DAPI. Scale bar, 10 μm. B. Wide-field microscopy images of stable, inducible cell lines expressing HABirA-tagged actin constructs as indicated, and stained with HA-antibody for construct expression, fluorescent streptavidin to visualize the biotinylated proteins and DAPI. Scale bar, 10 μm. C. Wide-field microscopy images of stable, inducible cell lines expressing Strep-HA or BirA*-tagged GFP used as controls in the MS analysis, stained with HA antibody and DAPI. Scale bar 10 μm.

**Supplementary figure 2 (S2).**
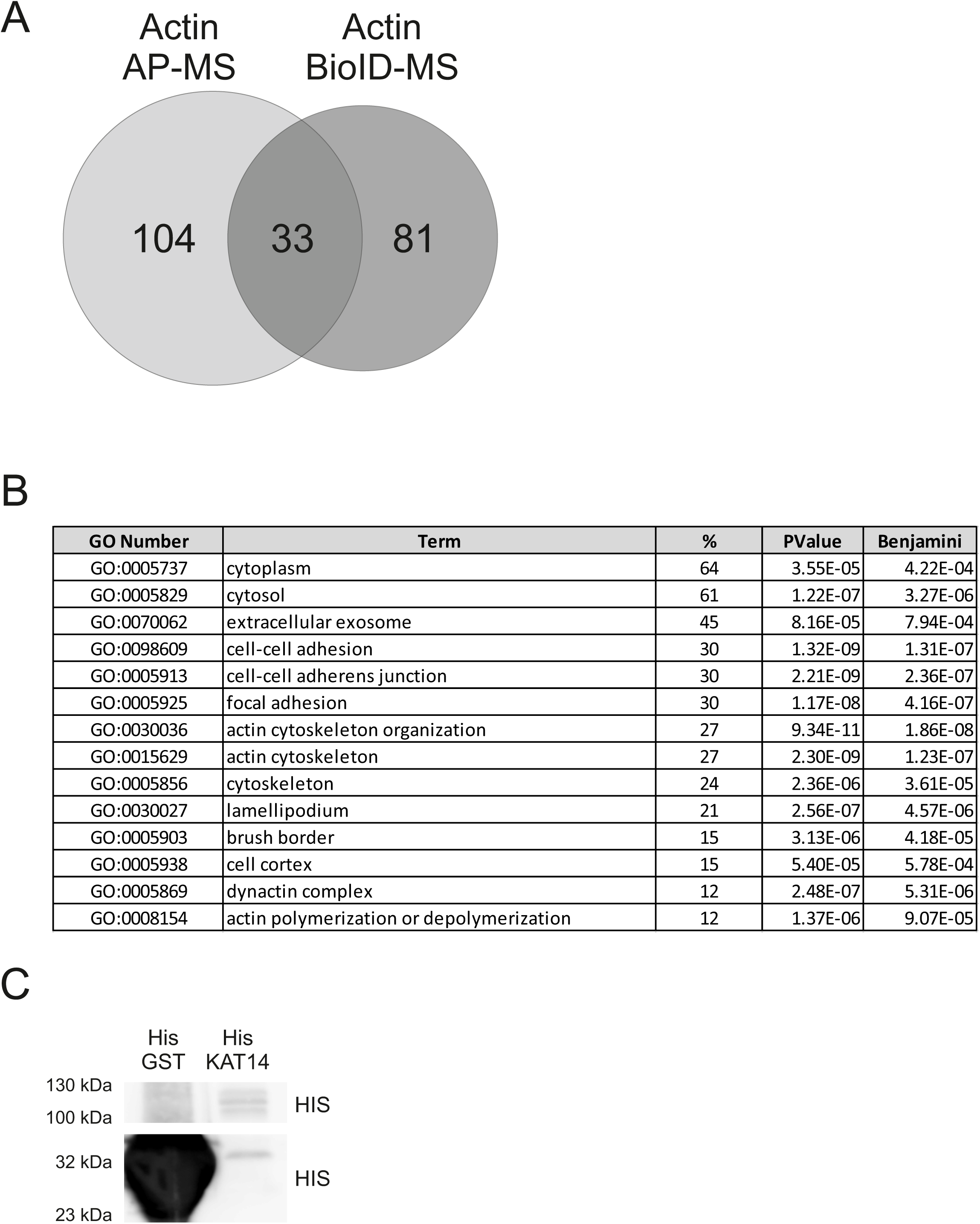

A. Number of unique and shared high confidence interactions from AP-MS and BioID screen for actin.

B. DAVID functional annotation chart (Cut off Benjamini 10^−4^) with GO Direct Biological Pathway (BP), Cellular Component (CC), Molecular Function (MF) from shared high confidence interactions from actin AP-MS and BioID screens hits.

C. Western blots of equal amounts of samples (33 %) loaded from the NTA Ni2+ pull down assay with His tagged GST and KAT14. HIS-HRP antibody was used to detect bound proteins in western blot. Related to figure 5D.

**Supplementary Table 1**

Raw data from the MS analysis

**Supplementary Table 2**

High-confidence interactions from the MS analysis

